# Intercellular CRISPR Screens Enhance the Discovery of Cancer Immunotherapy Targets

**DOI:** 10.1101/2022.03.03.482805

**Authors:** Soorin Yim, Woochang Hwang, Namshik Han, Doheon Lee

**Affiliations:** Department of Bio and Brain Engineering, KAIST, Daejeon, Republic of Korea; Bio-Synergy Research Center, Daejeon, Republic of Korea; Milner Therapeutics Institute, University of Cambridge, Cambridge, UK; Cambridge Centre for AI in Medicine, Department of Applied Mathematics and Theoretical Physics, University of Cambridge, Cambridge, UK

**Keywords:** intercellular interactions, ligand-receptor interactions, cell-cell communication, target discovery, immune checkpoint inhibitors, genome-wide CRISPR screen, triple-negative breast cancer, cytotoxic T cells

## Abstract

Cancer immunotherapy works through the interplay between immune and cancer cells. Particularly, interactions between cytotoxic T lymphocytes (CTLs) and cancer cells, such as *PDCD1* (PD-1) and *CD274* (PD-L1), are crucial for removing cancer cells. However, immune checkpoint inhibitors targeting these interactions are effective only to a subset of patients, requiring the development of novel immunotherapy drugs with novel targets.

Genome-wide clustered regularly interspaced short palindromic repeats (CRISPR) screening in either cancer or immune cells has been used to discover regulators of immune cell function as immunotherapeutic targets. However, the method has two main limitations. First, performing CRISPR screens in one cell type alone makes it difficult to identify essential intercellular interactions due to the focus on single genes instead of interactions. Second, pooled screening is associated with high noise levels. Therefore, we propose intercellular CRISPR screens, which perform genome-wide CRISPR screening in every interacting cell type to discover intercellular interactions as immunotherapeutic targets.

Intercellular CRISPR screens use two individual genome-wide CRISPR screens one each in immune and cancer cells to evaluate intercellular interactions that are crucial for killing cancer cells. We used two publicly available genome-wide CRISPR screening datasets obtained while triple-negative breast cancer (TNBC) cells and CTLs were interacting. We analyzed 4825 interactions between 1391 ligands and receptors on TNBC cells and CTLs to assess the effects of intercellular interactions on CTL function by incorporating both CRISPR datasets and the expression levels of ligands and receptors.

Our results showed that intercellular CRISPR screens discovered targets of approved drugs, a few of which were not identifiable using single datasets. To quantitatively evaluate the method’s performance, we used data for cytokines and costimulatory molecules because they constitute the majority of immunotherapeutic targets. Combining both CRISPR datasets improved the F1 score of discovering these genes relative to using single CRISPR datasets by more than twice.

Our results indicate that intercellular CRISPR screens can identify novel immune-oncology targets that were not obtained using individual CRISPR screens. The pipeline can be extended to other cancer and immune cell types, such as natural killer cells, to identify important intercellular interactions as potential immunotherapeutic targets.

## 1 Introduction

Cancer immunotherapy invigorates the immune system to fight cancer. This inevitably involves interactions between immune cells, and in particular cytotoxic T lymphocytes (CTLs), and cancer cells (1). For example, the well-known immune checkpoint inhibitor pembrolizumab is used to treat various types of cancer, including melanoma and triple-negative breast cancer (TNBC), as it targets *PDCD1* (PD-1) and interferes with *CD274* (PD-L1) interaction. The inhibitor prevents *CD274* from suppressing CTL function, enabling cancer cell removal (2). However, immune checkpoint inhibitors are effective in a subset of patients, with only 12.46% of cancer patients showing the benefits of treatment in the United States (3). To increase the number of patients who can benefit from the persistent effects of immunotherapy, novel immunotherapeutic drugs and targets are required.

Genome-wide clustered regularly interspaced short palindromic repeats (CRISPR) screens have been increasingly used to systemically discover targets for cancer therapy. Such efforts are especially evident for targeted therapy, where a large consortium called the Cancer Dependency Map performed 1076 genome-wide CRISPR screens in 908 cell lines (4, 5). Additionally, genome-wide CRISPR screening is being used for immunotherapy as well. For example, pooled CRISPR screening in immune or cancer cells has been used to evaluate immune cell function regulators and identify immunotherapeutic targets (6, 7). Dong et al. edited CTLs to identify which gene knockouts increased *in vivo* tumor infiltration and *in vitro* degranulation of CTLs (8). Further, Lawson et al. used CRISPR in cancer cells to identify regulators of CTL-mediated killing (9).

However, CRISPR screening analyses that aim to identify immunotherapeutic targets have three main limitations. First, most studies have not focused on intercellular interactions, which underlie the mechanisms for how immunotherapy works, and have instead focused on single genes. However, focusing on a single gene is insufficient as genes can interact with multiple partners (10). The drawbacks of this approach are evident when analyzing a ligand with opposing effects depending on the receptor it binds (11). For example, *CD80* can activate or suppress T cells upon binding to *CD28* and *CTLA4*, respectively (12). Such competition was also observed between *PVR, CD226*, and *TIGIT* (**Figure 1A**) (13). Moreover, our analysis showed that a large proportion of immuno-oncology (IO) drugs and their targets are derived from intercellular communication molecules such as adhesion molecules, surface antigens, and membrane receptors (**Figure 1B, C, and Supplementary Table 1**). Intercellular interactions are more potent drug targets than cytosolic proteins because they are exposed from the cellular membrane and are more druggable (14). Second, a few studies have performed CRISPR screening using monocultured cells, even though monocultures do not reflect cell-cell communication. Studies that have used CRISPR screening under co-culture or other settings that can reflect intercellular interactions may also be limiting because the screening was performed in one cell type alone, i.e., in immune or cancer cells. Performing CRISPR screening in one cell type makes it difficult to pinpoint essential intercellular interactions due to the focus on a particular gene rather than analyzing interactions (15). Lastly, genome-wide CRISPR screens inevitably entail high technical noise owing to their high throughput (16). To overcome these limitations and robustly discover essential intercellular interactions, CRISPR screens should be performed in every cell type that interacts with each other.

**Figure 1.**
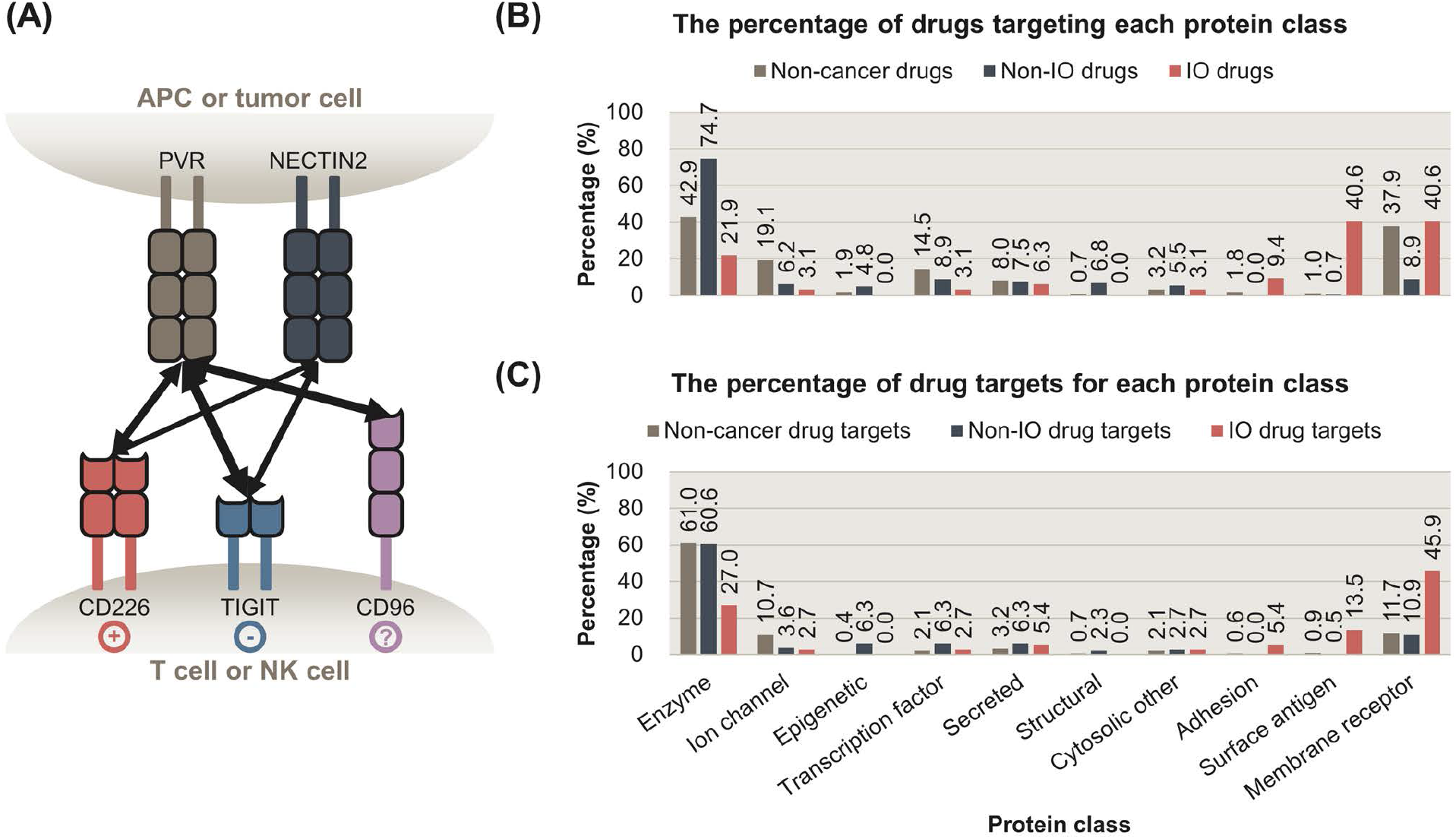
Intercellular interactions are potent targets for cancer immunotherapy. **(A)** Intercellular interactions between *PVR, NECTIN2, CD226, TIGIT*, and *CD96. PVR* and *NECTIN2* are expressed on antigen-presenting cells (APCs) and some tumor cells. Their receptors, *CD226, TIGIT*, and *CD96* are expressed on T cells or natural killer cells (NK cells). Upon binding to *PVR* or *NECTIN2, CD226* and *TIGIT* trigger stimulatory and inhibitory signals, respectively. Whether the binding of *PVR* to *CD96* delivers stimulatory or inhibitory signals is to be determined. The multiplicity of these interactions highlights the importance of focusing on intercellular interactions, rather than a single ligand or a receptor. **(B)** The percentage of non-cancer and cancer drugs whose targets belong to each protein class. Cancer drugs were further categorized into non-immuno-oncology (non-IO) and IO drugs. **(C)** The percentage of proteins targeted by non-cancer, non-IO, and IO drugs that belong to each protein class. Membrane receptors, surface antigens, and adhesion proteins are preferentially targeted by IO drugs, whereas enzymes are less favorably targeted by IO drugs. APC: antigen-presenting cell, NK cell: natural killer cell, IO: immuno-oncology.

In this study, we suggest intercellular CRISPR screens and a corresponding analysis method to evaluate intercellular interactions as IO targets. Intercellular CRISPR screens involve two genome-wide CRISPR screens, one each in immune and cancer cells. Both screenings occur during the interaction between the two cell types to identify critical intercellular interactions that kill cancer cells. Through combining both datasets, we calculated the ‘intercellular normZ score’ for each intercellular interaction to quantify its potential as an immunotherapeutic target. As a proof of concept, we combined two publicly available genome-wide CRISPR screening datasets obtained while cancer cells and CTLs were interacting with each other (8, 9). Our results showed that intercellular CRISPR screens are more effective in identifying IO targets than CRISPR screens performed alone.

## 2 Materials and Methods

### 2.1 Classification of Cancer Drugs and Their Targets

#### 2.1.1 Collection of Approved Cancer Drugs from DrugBank

The list of drugs, along with their development status (‘groups’), anatomical therapeutic chemical (ATC) codes, and target information was collected from DrugBank 5.1.8 (17). Among 14,585 drugs in DrugBank, 4207 belonged to the approved group, indicating that the drugs were approved in at least one jurisdiction at some time point. We set two criteria, either of which should be met for a drug to be termed a cancer drug; (1) The drugs should have ATC codes starting with ‘L01’ (antineoplastic agents), or (2) drugs belonging to the ‘Cancer immunotherapy’ category (accession number DBCAT005215) that were used in cancer treatment owing to their immunological effects. The DrugBank Clinical API was used to retrieve 72 approved drugs that belong to the ‘Cancer immunotherapy’ category. Altogether 221 cancer drugs were identified.

#### 2.1.2 Classification of Cancer Drugs into IO vs Non-IO Using DrugBank and IO Landscape

Two resources were used to classify cancer drugs as IO and non-IO drugs. The first resource was the ‘Cancer immunotherapy’ category (accession number DBCAT005215) from DrugBank. The second was the IO Landscape, developed by the Cancer Research Institute to catalog the developmental status of IO drugs (18).

Among the ‘Cancer immunotherapy’ drugs from DrugBank, a few drugs such as trastuzumab, a *ERBB2* (HER2) inhibitor, are monoclonal antibodies that can be regarded as a targeted therapy rather than immunotherapy (19). However, a few monoclonal antibodies including pembrolizumab, a *PDCD1* inhibitor, are used as immunotherapeutic drugs. Therefore, we set two criteria, one of which should be met to classify a drug as an IO drug; (1) The drug belongs to the DrugBank ‘Cancer immunotherapy’ category and does not have ATC codes starting with ‘L01XC’ (monoclonal antibodies), or (2) A drug belongs to the ‘Cancer immunotherapy’ category, has ATC codes starting with ‘L01XC’, and is listed on IO Landscape. As a result, 47 cancer drugs were classified as IO drugs. The remaining 174 cancer drugs were classified as non-IO drugs.

#### 2.1.3 Protein Target Classification using ChEMBL

Drug target information was collected from DrugBank. We used the classification term ‘targets’ and excluded ‘enzymes’, ‘transporters’, and ‘carriers’. As a result, 282 targets for 185 approved cancer drugs were identified.

The targets were classified using ChEMBL 29 (20). We used Level 1 classification terms to classify proteins into 14 categories. If a protein was classified as a “child” term, the term was mapped to the corresponding Level 1 terms based on the classification hierarchy. A target can have more than one category. For example, B-cell receptor CD22 was classified into the ‘Adhesion’, ‘Membrane receptor’, and ‘Surface antigen’ categories.

Drugs were categorized based on their target classification. If a drug had multiple targets, it was considered to belong to all classes into which its targets were classified.

### 2.2 Intercellular CRISPR Screens: Integration of CRISPR Screens in Immune and Cancer Cells to Prioritize Intercellular Interactions for IO Target Discovery

#### 2.2.1 Data Sources

##### 2.2.1.1 Genome-wide Pooled CRISPR Screens in CTLs

Genome-wide pooled CRISPR screens of CTLs were obtained from Dong et al. (8). The screen aimed to identify genes whose knockout increased the infiltration of CTLs into the tumor tissue *in vivo*. To achieve this goal, CTLs were transduced with a mouse genome-scale single guide RNA (sgRNA) library and were subsequently injected into *Rag1-/-* mice with TNBC, E0771 tumors (**Figure 2A**), and tumor tissues were harvested. The harvested tumor samples were compared with cellular libraries of infected CTLs that served as control samples to identify the enriched sgRNAs and genes in the tumor samples.

**Figure 2.**
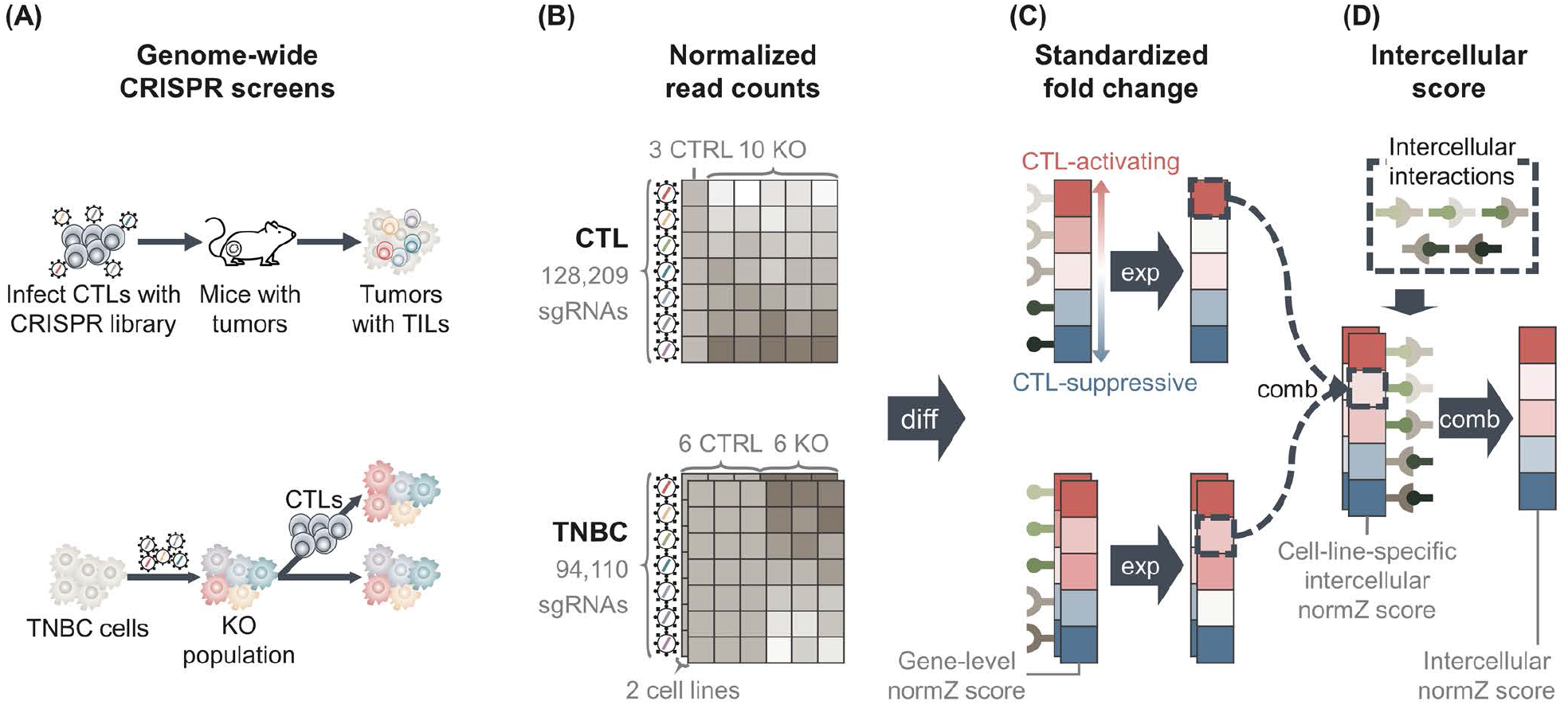
Methods overview. **(A)** Data from two genome-wide pooled clustered regularly interspaced short palindromic repeats (CRISPR) screens were used. One CRISPR screen edited cytotoxic T lymphocytes (CTLs) to identify genes whose knockout increased the infiltration of CTLs into tumor tissue (upper) (8). The second screen edited two triple-negative breast cancer (TNBC) cell lines to identify genes whose knockout regulates the evasion of TNBC cells from CTL-mediated killing (lower) (9). **(B)** Genome-wide CRISPR screens yielded normalized read count matrices showing the amount of sgRNAs in each sample. TNBC CRISPR screen data had two matrices, one for each TNBC cell line. **(C)** We performed differential analysis to calculate fold changes of genes between knockout and control samples, yielding ‘Gene-level normZ scores’. Positive scores were assigned genes that were more likely to activate CTL function and vice versa. The scores of genes that were not expressed in CTLs/TNBC cells lines were set to zero. **(D)** We collected information for intercellular interactions from public databases and calculated the score of each intercellular interaction by ‘combining’ gene-level normZ scores of the interactants. Because two TNBC cell lines were used, two ‘cell-line specific intercellular normZ scores’ were obtained, one for each TNBC cell line. We combined cell-line-specific intercellular normZ scores to obtain the final intercellular normZ score. TIL: tumor-infiltrating lymphocyte, KO: knockout, CTRL: control, diff: differential analysis, exp: expression, comb: combination.

The normalized read count matrix was downloaded from Table S1 (8). The dataset contained the normalized read counts of 128,209 sgRNAs targeting 21,786 genes from three cellular libraries and ten knockout samples (**Figure 2B**).

##### 2.2.1.2 Gene expression in CTLs

In the study by Dong et al. (8), single-cell RNA sequencing was used to identify genes expressed in CD8+ tumor-infiltrating lymphocytes. From the paper, Table S2 was used to obtain a list of genes expressed in CTLs as follows: First, low-quality cells with small library sizes (the number of unique molecule identifiers ≥4 standard deviations below the mean) or diversities (the number of detected genes ≥4 standard deviations below the mean), or large mitochondrial genes (the proportion of mitochondrial genes ≥4 standard deviations above the mean) were removed. Genes with low variance were regarded as being unexpressed. After converting Ensemble IDs into gene symbols, 7874 genes were identified expressed in CTLs (21).

##### 2.2.1.3 Genome-wide Pooled CRISPR Screens in TNBC Cell Lines

A genome-wide pooled CRISPR screen in two mouse TNBC cell lines, 4T1 and EMT6, was conducted by Lawson et al. (9). The screening was performed to identify cancer genes whose knockout regulates the evasion of CTL-mediated killing of cancer cells. To this end, cancer cells were infected with the mouse Toronto knockout library and co-cultured with activated CTLs (**Figure 2A**). The number of sgRNAs in co-cultured cancer cells was compared with that in monocultured cancer cells as control samples. We downloaded two normalized read count matrices consisting of 94,110 sgRNAs targeting 19,459 genes in six co-culture and six monoculture samples for each cell line from Supplementary Table 2 (9) (**Figure 2B**).

##### 2.2.1.4 Gene expression in TNBC Cell Lines

To identify genes that were expressed in the mouse TNBC cell lines 4T1 and EMT6, bulk RNA sequencing data were obtained from Supplementary Table 6 (9), from which the TNBC CRISPR screen data was collected. Two replicates were used for each TNBC cell line. A gene was considered as being expressed in a cell line if the mean expression level across replicates was at least ten fragments per kilobase of exon per million (FPKM) (22). If a gene had multiple probes, the probe with the highest average expression level was used. As a result, 6411 and 6518 genes were identified in 4T1 and EMT6, respectively.

##### 2.2.1.5 Intercellular Interactions from ConnectomeDB2020 and CellPhoneDB

Intercellular interaction data were collected from two sources: ConnectomeDB2020 (23) and CellPhoneDB (24). ConnectomeDB2020 contains 2293 manually curated ligand-receptor interactions. CellPhoneDB is a database of ligands, receptors, and describes 1396 interactions between these. Although both databases provide information on interaction data from humans, the CRISPR screens were performed in mouse cells. Therefore, we used orthologs to obtain potential ligand-receptor interactions in mice (21, 23). As a result, 4825 intercellular interactions between 1391 ligands and receptors were identified.

#### 2.2.2 Differential Analysis to Calculate Fold Change of Genes

We performed differential analysis using the drugZ algorithm (25) to identify genes that were significantly differentially enriched in knockout samples compared with control samples in each CRISPR dataset (**Figure 2C**). First, the drugZ algorithm compares sgRNA counts in knockout samples against control samples to calculate the fold changes in the amount of sgRNAs. Next, the algorithm standardizes the fold change by dividing it by the standard deviation and converts it into a *z*-score. To robustly estimate the standard deviation, drugZ uses empirical Bayes methods by borrowing information from sgRNAs with similar read counts in the control samples. Next, it combines the *z*-scores of individual sgRNAs targeting the same gene by summing the individual *z*-scores and dividing the sum by the square root of the number of summed sgRNAs. This yields the final normZ score, which follows a standard normal distribution.

For the TNBC CRISPR screening dataset, we used drugZ algorithm to calculate fold changes of genes co-cultured with CTLs compared to monocultured TNBC cells. As a result, genes whose knockout help TNBC cells escape from CTL-mediated killing obtained positive normZ scores. Since knockouts of these genes prevent CTLs from killing TNBC cells, they may be components of CTL-mediated cancer cell removal. For the CTL CRISPR screening dataset, we calculated the fold changes of genes in cell libraries against the tumor samples in order to give negative scores to CTL-suppressive genes. Genes whose knockout increased CTL infiltration into tumor tissue were enriched in the tumor sample, and therefore given negative scores. These genes may prevent CTL infiltration in their normal state. As a result, positive normZ scores were given to the genes that activate CTL function, while CTL-suppressive genes had negative normZ scores for both CRISPR screening datasets.

#### 2.2.3 Calculation of Intercellular normZ Score for Each Interaction

We combined two genome-wide CRISPR screening datasets to calculate the intercellular normZ scores. Before combining the CTL CRISPR and TNBC CRISPR screens, we set the normZ scores of unexpressed genes to zero for each cell type to reduce noise resulting from the high-throughput pooled screening procedure (5) (**Figure 2C**).

After zero out the normZ scores of unexpressed genes for each cell type, the normZ score of a ligand from one cell type and the corresponding receptor’s normZ score from the other cell type were summed to calculate the cell-line-specific intercellular normZ score (**Figure 2D, Supplementary Figure 1**). Because the gene-level normZ scores follow a standard normal distribution, we divided the sum by square root of two, i.e. the number of interacting genes, to obtain intercellular normZ scores that follow a standard normal distribution. As a result, we obtained two intercellular normZ scores, one each from 4T1-CTL and EMT6-CTL. For each intercellular interaction, we summed both normZ scores and divided the sum by the square root of two, i.e., the number of CTL-TNBC pairs, to obtain the final intercellular normZ score (**Figure 2D**). Because normZ scores follow the standard normal distribution (*z*-scores), the corresponding *p*-values and false discovery rates (FDRs) were calculated based on the normZ scores (25) (**Supplementary Table 2**).

#### 2.2.4 Alternative Methods to Calculate Intercellular normZ Scores

We devised two alternative methods to calculate the intercellular normZ scores. The first is ‘Strict’, which explicitly requires normZ scores to have the same sign when summed. If two normZ scores with different signs have to be summed, the ‘Strict’ method sets the result as zero instead of summing the scores. On the other hand, the proposed method in 2.2.3 does not put any constraints on the signs when two normZ scores are added, hence we named it as ‘Tolerant’. The second alternative method is ‘Composite’, a hybrid of ‘Strict’ and ‘Tolerant’. When the gene-level normZ scores of CTL and TNBC screens are summed, ‘Composite’ requires them to have the same sign. However, the intercellular normZ scores for 4T1-CTL and EMT6-CTL are not required to have the same sign to calculate the final score because cancer cell lines are highly heterogenous and they may not show the same immunological effect.

### 2.3 Collection of Well-Known Immunomodulators for Performance Evaluation

#### 2.3.1 Gold-Standard Dataset: Targets of Approved Drugs and Phase III Clinical Trial Drug Candidates for Immunotherapy

To benchmark the performance of intercellular CRISPR screens over using either of the screening data alone, a gold-standard dataset was obtained. Approved immunotherapy drugs that modulate CTLs were used as the gold-standard dataset to discover novel immunotherapeutic targets using CTL-related CRISPR screening data. We downloaded IO Landscape data and filtered drugs whose clinical stage was ‘Approved’ (18). Among 141 approved drugs, 97 drugs were classified as ‘T-cell targeted immunomodulators’ or ‘Other immunomodulators’. Among these, we retrieved 32 drugs whose target cell types were ‘APC/T cell’ or ‘T cell’. We manually inspected the 32 drugs and finalized the list of gold-standard drugs. However, only ten intercellular interactions were targeted by the approved drugs. Therefore, we obtained information for other drug candidates that modulate the function of CTLs and whose clinical stages were ‘Phase III’. As a result, 38 intercellular interactions between 47 genes and their effects on CTL function were identified (**Supplementary Table 3**).

#### 2.3.2 Silver Standard Dataset: Cytokines and Co-stimulatory Molecules

The size of the gold-standard dataset was still limited to quantitatively evaluating and comparing the methods. Therefore, we collected a silver standard dataset composed of well-known immunomodulators as potential immunotherapeutic targets. These immunomodulators include cytokines, and co-stimulatory and co-inhibitory molecules since they compose the majority of targets of T cell modulators and other modulators (26). Because we used CRISPR datasets obtained while cancer cells and CTLs were interacting, we collected CTL-related immunomodulators only (12, 13, 27-44). As a result, we obtained 79 and 30 intercellular interactions known to activate and suppress CTL function, respectively.

### 2.4 Performance Comparison

#### 2.4.1 Aggregation of Intercellular normZ Scores into Gene-Level Scores

Intercellular CRISPR screen evaluates each interaction, whereas CTL CRISPR and TNBC CRISPR screens evaluate each gene. To compare the intercellular CRISPR screen with the CTL and TNBC CRISPR screens, we aggregated interaction-level intercellular normZ scores into gene-level scores (**Supplementary Figure 2**). We enumerated all intercellular interactions involving each gene in CTL/TNBC cells. We hypothesized that strong interactions would result in intercellular normZ scores with high absolute values. Therefore, we used the intercellular normZ score with the highest absolute value as the score for the gene in the corresponding cell type.

#### 2.4.2 Performance Evaluation Metrics

We used the area under the receiver operating characteristic curve (AUROC), precision, recall, and F1 scores to evaluate the ability of CRISPR screens to discover the silver standard dataset. For the AUROC, we used the normZ scores. For precision, recall, and F1 score, we used FDR < 5% to classify normZ scores as CTL-activating, CTL-suppressive, and unknown. Genes/interactions with FDR ≥ 5% were classified as unknown. The remaining genes/interactions with positive and negative normZ scores were classified as CTL-activating and CTL-suppressive, respectively.

Because we have three classes, CTL-activating, CTL-suppressive, and unknown, we used two averaging schemes that are widely used in multiclass classifications. The first scheme is micro-averaging, which weighs each instance equally. On the other hand, macro-averaging regards each class as equally important, giving more weight to instances from minor classes.

## 3 Results

### 3.1 Intercellular CRISPR Screens Identify Approved Targets for Cancer Immunotherapy

To discover novel immunotherapeutic targets, we focused on intercellular interactions instead of single genes. We propose the use of intercellular CRISPR screens as a pipeline to discover potentially therapeutic interactions between immune and cancer cells. As a proof of concept, we used an intercellular CRISPR screen to identify the interactions that affect CTLs in TNBC (**Figure 2**). We used two genome-wide pooled CRISPR screen datasets that were collected while CTLs were interacting with TNBC cells (8, 9). Next, we collected intercellular interactions from CellPhoneDB and ConnectomeDB2020 (23, 24). By combining the two CRISPR screen datasets and the expression level of each gene in CTLs and TNBC cell lines, we quantified the extent to which each intercellular interaction affected CTL function.

**Figure 3A** shows that the intercellular CRISPR screen (‘Tolerant’) can identify both CTL-activating and suppressive interactions which can increase or inhibit CTL-mediated cancer cell removal, respectively. In contrast, the CTL and TNBC CRISPR screens are more suited for identifying CTL-suppressive and activating genes, respectively. The results can be explained as follows: knockout of CTL-activating genes decreases the number of CTLs, resulting in increased TNBC cell numbers. In contrast, the knockout of CTL-suppressive genes increases the number of CTLs, resulting in decreased TNBC cells. CTL CRISPR screens are more suitable for identifying genes that increase the number of CTLs, i.e., CTL-suppressive genes, whereas TNBC CRISPR screens are more suitable for identifying CTL-activating genes which can increase TNBC cell numbers. By combining both CRISPR screens, intercellular CRISPR screens can effectively identify CTL-activating and suppressive genes.

**Figure 3.**
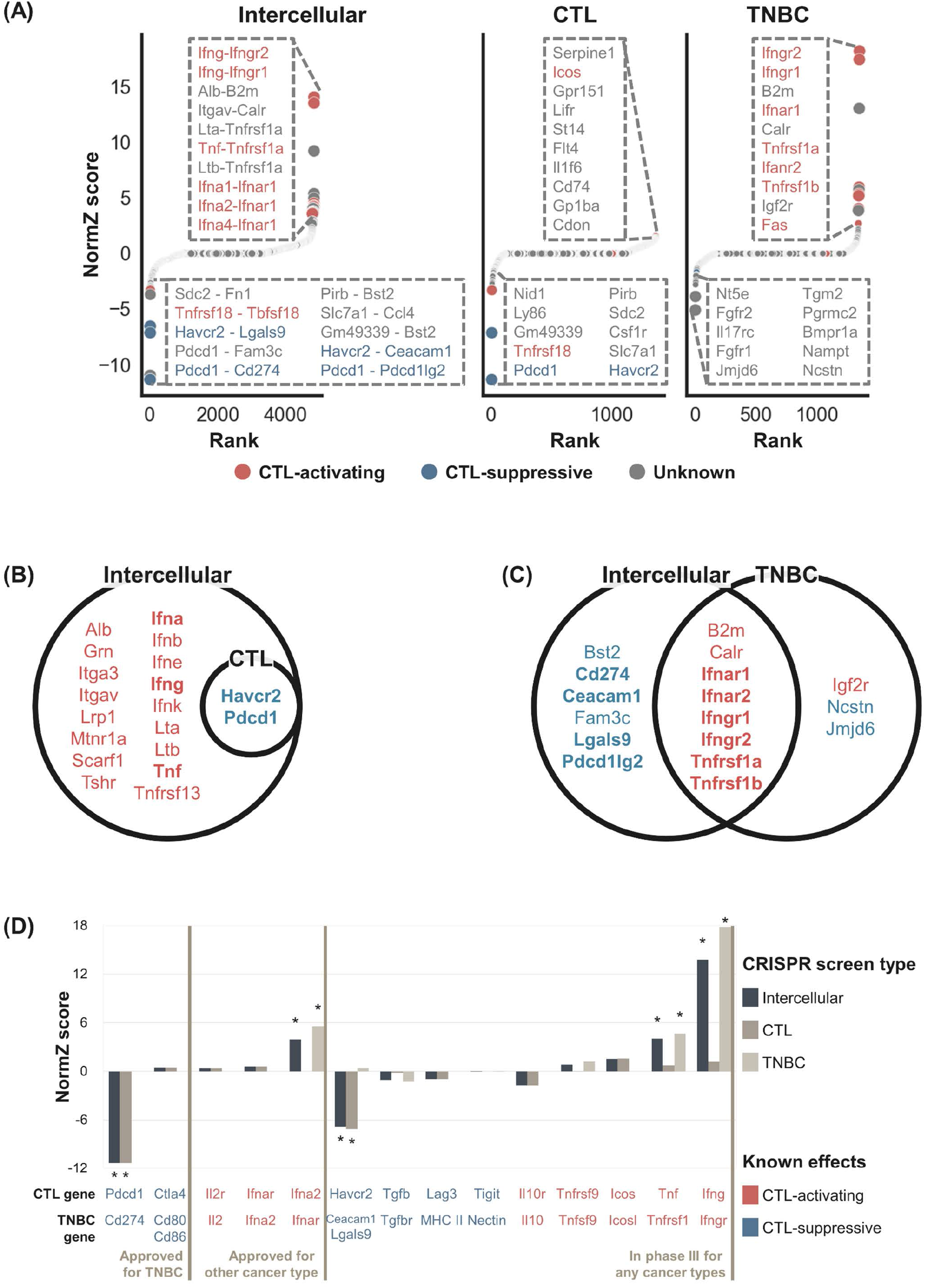
NormZ scores from the intercellular, CTL, and TNBC CRISPR screens. **(A)** Rank-ordered normZ scores from the intercellular (left), CTL (middle), and TNBC (right) CRISPR screens. Interactions/genes known to activate and suppress CTL function are marked in red and blue, respectively. The dot sizes are negatively proportional to the FDR. The top ten interactions/genes are represented in the inset. **(B)** Statistically significant (FDR < 5%) genes from the gene-level intercellular and CTL CRISPR screens. **(C)** Statistically significant (FDR < 5%) genes from the gene-level intercellular and TNBC CRISPR screens. Genes with positive and negative normZ scores are marked with red and blue, respectively. Well-known immunomodulators from the silver standard data are marked in bold. **(D)** The intercellular, CTL, and TNBC normZ scores of interactions/genes targeted by approved immunotherapeutic drugs, or phase III clinical trial drug candidates. * Statistically significant (FDR < 5%) interactions/genes.

The results for gene-level comparisons showed a similar tendency (**Figure 3B, C**). The CTL CRISPR screen identified CTL-suppressive genes whereas genes discovered from TNBC CRISPR screens were biased towards CTL-activating ones. In contrast, intercellular CRISPR screens can help identify both CTL-activating and suppressive genes.

To benchmark the use of intercellular CRISPR screens relative to the use of either CRISPR screen alone, we obtained a gold-standard dataset containing data for 38 intercellular interactions targeted by approved IO drugs or phase III clinical trial drug candidates for immunotherapy. **Figure 3D** shows that CTL-suppressing and activating interactions tended to have negative and positive scores, respectively. Significant interactions were mostly related to cancer antigen presentation and killing of cancer cells, reflecting the experimental endpoint of the used CRISPR screen (45). This may be the reason why *Ctla4* obtained insignificant scores since it is related to the proliferation of CTLs in lymph nodes rather than killing cancer cells in peripheral tissue (45, 46). A few interactions were missed when CRISPR screen data were used alone. For example, the TNBC CRISPR screen failed to identify *Cd274* (Pd-l1) because it was not expressed in either 4T1 or EMT6 cells. However, this result is biologically plausible because not all patients express *PD-L1* or respond to anti-*PD-L1* therapy (47). In contrast, the CTL CRISPR dataset failed to identify interferons alpha and gamma.

These results indicated that intercellular CRISPR screens combined the complementary CRISPR datasets to identify IO targets more comprehensively. However, a few gold-standard interactions had negligible scores. For example, aldesleukin is a recombinant interleukin-2 that has been approved for the treatment of renal cell carcinoma. However, the normZ scores of *IL2* and *IL2R* were close to zero in the CTL and TNBC CRISPR screens, respectively. Interestingly, we found that interleukin-2 failed to demonstrate its efficacy in breast cancer patients during a phase III clinical trial (48). Similarly, pegylated recombinant interleukin-10 or pegilodecakin had low efficacy against pancreatic ductal adenocarcinoma during a phase III clinical trial (49).

In summary, the intercellular CRISPR screen identified two intercellular interactions, *Pdcd1*-*Cd274* and *Ifna2*-*Ifnar* targeted by approved IO drugs pembrolizumab and peginterferon alfa-2b, respectively. In addition, our results suggested *Havcr2*-*Lgals9* and *Ifng*-*Ifngr*, targeted by sabatolimab and interferon gamma-1b respectively, as potentially therapeutic interactions. These results indicate that intercellular CRISPR screens can be used to discover effective targets during the early drug development stages.

### 3.2 Intercellular CRISPR Screens Outperform Single CRISPR Screens

We performed three quantitative evaluations to compare the performances of different CRISPR screens. First, we evaluated the performance of intercellular CRISPR screens calculated by the ‘Tolerant’ method. Next, we compared the ‘Tolerant’ method with two alternative methods, ‘Strict’ and ‘Composite’, to identify the best way to calculate intercellular normZ scores. Lastly, we compared the performance of intercellular CRISPR screens with CTL and TNBC CRISPR screens to estimate the degree of performance enhancement. Because there are few IO drugs, the gold-standard dataset may not be appropriate for the quantitative evaluation. Therefore, we used a silver standard dataset containing cytokines and co-stimulatory molecules as potential immunotherapeutic targets.

Cytokines and co-stimulatory molecules were selected since they compose the majority of targets of immunomodulatory drugs (26). Because intercellular normZ scores are continuous, they were binarized based on an FDR < 5% to make predictions.

The confusion matrix of the ‘Composite’ method is shown in **Figure 4A**. Among 79 CTL-activating interactions in the silver standard dataset, 30 were correctly classified as CTL-activating, whereas the remaining 49 were classified as unknown. Among 30 CTL-suppressive interactions, four were classified as CTL-suppressive, whereas the remaining 26 were classified as unknown. Even though 21 CTL-suppressive interactions obtained negative normZ scores, only four were statistically significant, i.e. FDR < 5%. Among 4716 interactions whose effects are unknown, 20 and one interaction were classified as CTL-activating and CTL-suppressive, respectively, and suggested as potential immunotherapeutic targets. Based on the confusion matrix, we evaluated the precision, recall, and F1 scores of intercellular CRISPR screen against the silver standard dataset. We used micro and macro-averaging to deal with multiclass classification. Micro-averaged precision, recall, and F1 score were 0.62, 0.31, 0.41, respectively (**Figure 4B**). The recall was lower than precision since we used a strict threshold, FDR < 5%, for the classification to identify highly confident potential targets.

**Figure 4.**
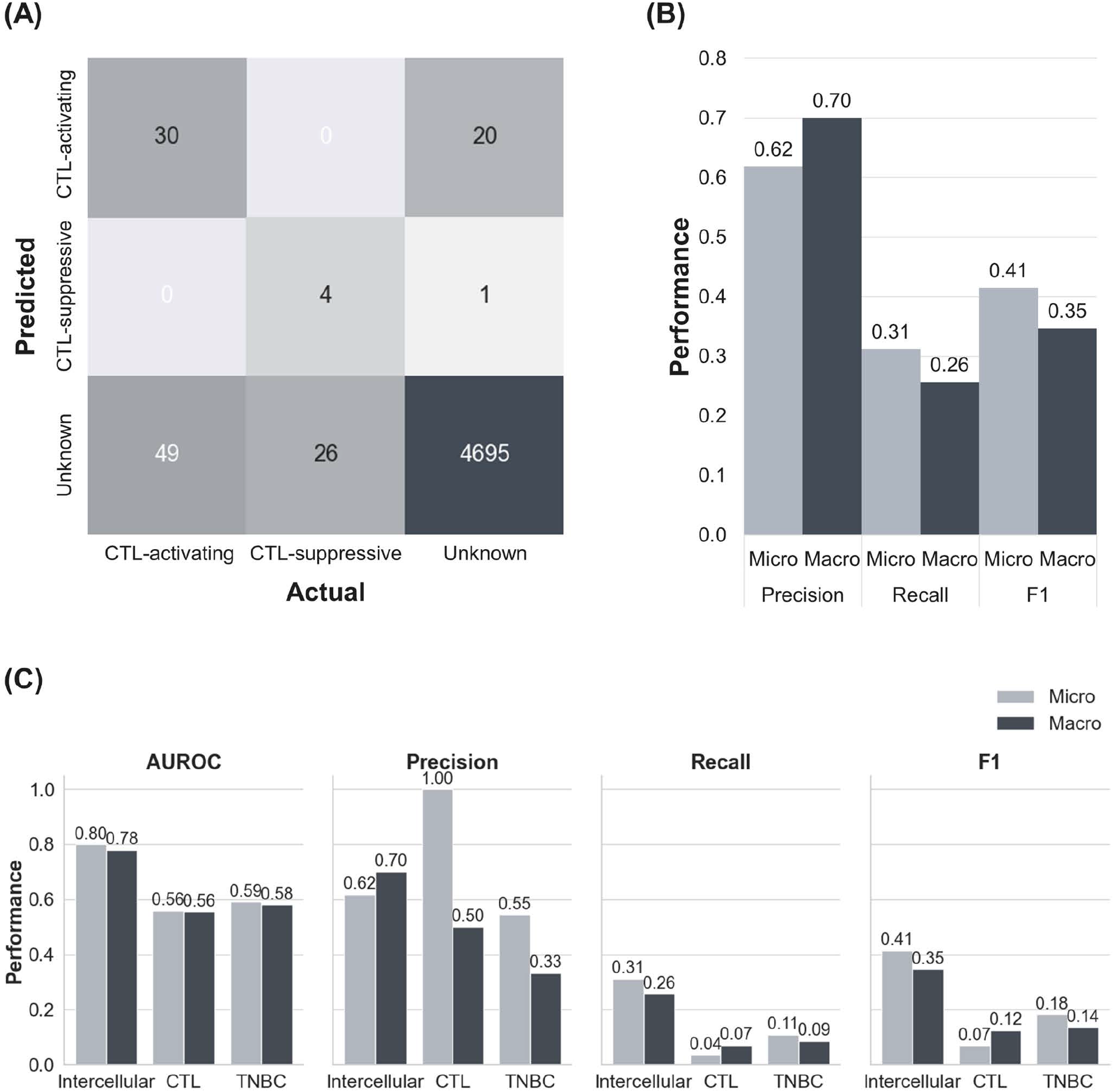
The performance of the intercellular CRISPR screen. (A) Confusion matrix of intercellular CRISPR screens. Predictions were made based on an FDR < 5%. (B) Precision, recall, and F1 scores of intercellular CRISPR screens. (C) AUROC, precision, recall, and F1 scores of CTL, TNBC, and intercellular CRISPR screens. AUROC: area under the receiver operating characteristic curve.

We evaluated two alternative methods to calculate the intercellular normZ scores (**Supplementary Figure 3**). The first method is ‘Strict’, which explicitly requires normZ scores to have the same sign when summed. We named the method in **Figure 2** as ‘Tolerant’ since it does not put any constraints on the signs of normZ scores. The second alternative method is ‘Composite’, which requires the gene-level normZ scores of CTL and TNBC screens to have the same sign. However, the intercellular normZ scores for 4T1-CTL and EMT6-CTL are not required to have the same sign under the ‘Composite’ method. We evaluated ‘Tolerant’, ‘Composite’, and ‘Strict’ when used with and without gene expression data. The best performance was obtained using the ‘Tolerant’ method with expression data, whereas the ‘Strict’ method with expression data performed the worst. For the ‘Composite’ and ‘Strict’ methods, the use of gene expression data reduced the performances due to the strict constraint on the normZ scores which resulted in many zeros. In contrast, including expression data improved the performance of the ‘Tolerant’ method. Therefore, we used the ‘Tolerant’ method with expression data to calculate the intercellular normZ scores.

Next, we compared the performance of the intercellular CRISPR screen relative to the CTL and TNBC CRISPR screens. We calculated the AUROC, precision, recall, and F1 scores for each CRISPR screen (**Figure 4C, Supplementary Figure 4**). The results showed that the intercellular CRISPR screen outperformed both individual CRISPR screens for all evaluation metrics except the precision of the CTL screen. However, the CTL screen identified two interactions alone, resulting in poor recall and F1 scores. In fact, the intercellular CRISPR screen improved the F1 score over single CRISPR screens more than twice. These results demonstrate that intercellular CRISPR screens can use complementary CRISPR screens to reduce the noise from high-throughput CRISPR screens to robustly identify immunotherapeutic targets.

### 3.3 Intercellular CRISPR Screens Identify Potential IO Targets

Based on the intercellular normZ scores, we identified potential IO targets with previously unknown effects. Among the 59 intercellular interactions whose absolute scores were ≥3, we finalized seven interactions based on the following criteria; (1) Interactions whose effects from CTL and TNBC CRISPR screen were concordant, and (2) interactions involving genes expressed in CTLs. We included genes that were not expressed in TNBC cell lines because a few genes, including *CD274*, may be expressed in only a subset of TNBC cell lines.

Among the seven interactions identified, five were partially supported by literature (**Table 1**). Three interactions involved *Tnfrsf1a/b*, which activate CTL function (41). In particular, *Tnfrsf1b*-mediated activation of CTLs showed tumor regression in syngeneic EMT6 models (50). In addition, a recent study reported that *Slc7a1* reduces memory T cells by activating mTORC1 (51), and *Sdc2* facilitates the removal of the T-cell receptor/CD3 complex from the cell membrane (52).

**Table 1.**
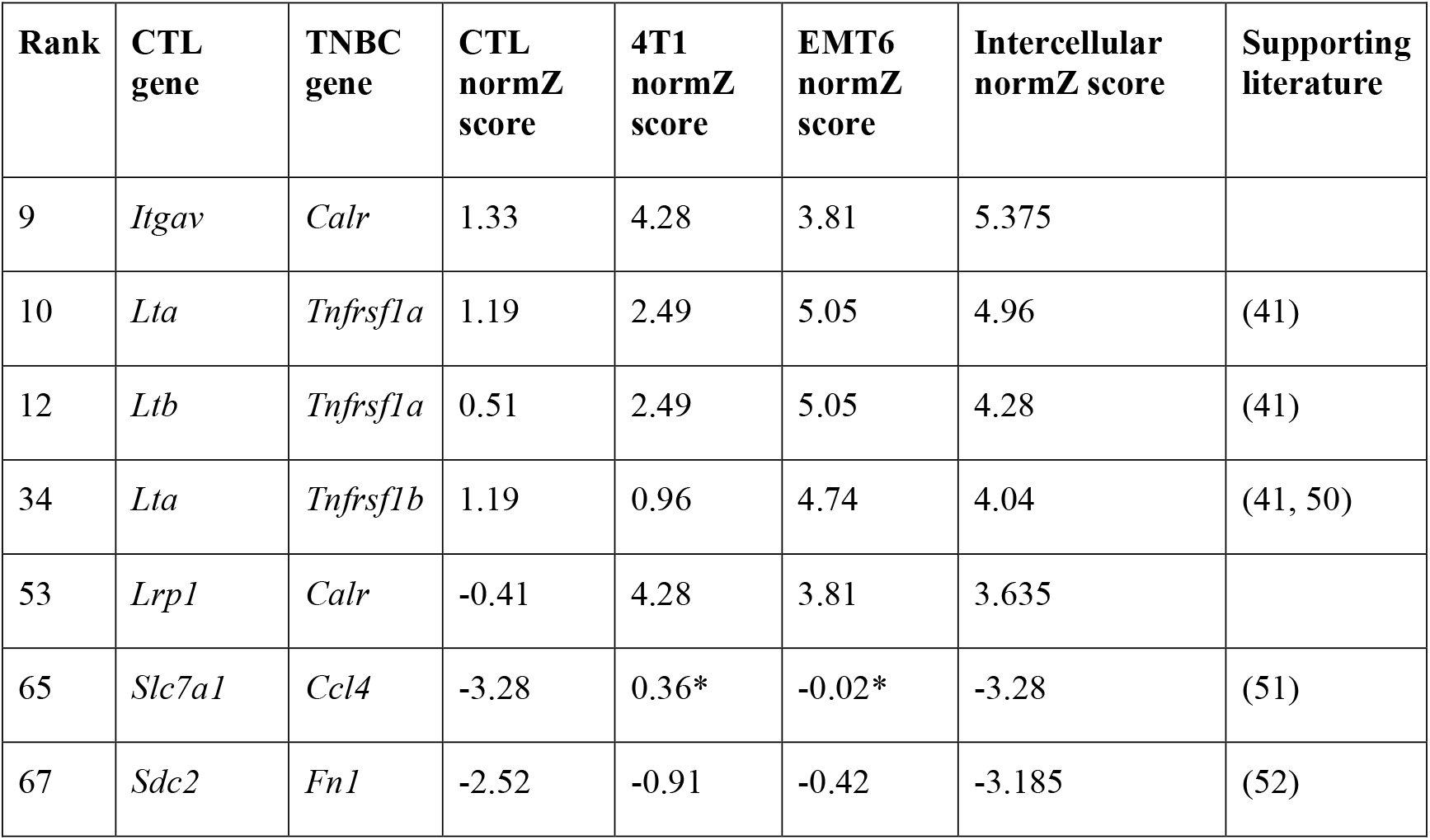
Highly ranked intercellular interactions for immunotherapeutic targets. * *Ccl4* was not expressed in 4T1 and EMT6 TNBC cell lines. All the other genes were expressed in the corresponding cell types.

In summary, our results suggested seven intercellular interactions as immunotherapeutic targets for TNBC. Among these, four may activate CTL function, for which agonists can be investigated as IO drugs. The remaining three were suggested to suppress CTL function, for which antagonists may be investigated to treat TNBC.

## 4 Discussion

In this study, we have shown the utility of intercellular CRISPR screens to discover interactions between immune and cancer cells as IO targets, instead of focusing on single genes. Intercellular CRISPR screens integrate two CRISPR screens, performed one each in immune and cancer cells, which are obtained when both cells interact with each other. We used an intercellular CRISPR screen to evaluate essential interactions between CTLs and TNBC cells and identified IO targets. Our results showed that CTL and TNBC CRISPR screens were complementary and the intercellular CRISPR screen outperformed the individual screens.

Although our method successfully identified approved IO targets and known immunomodulators, it has two limitations. First, the experimental settings for the CTL and TNBC CRISPR screens were different. The CTL and TNBC CRISPR screens were used *in vivo* and *in vitro*, respectively. In addition, the screens used different TNBC cell lines. Although we tried to find datasets with similar experimental conditions, there are few studies of genome-wide CRISPR screens using the same immune cell types and cancer cells. Second, public databases were used to identify putative interactions between CTLs and TNBC cells. Growing efforts have been made to infer cell-type-specific intercellular interactions by combining information from public databases with large-scale gene expression data (53). Although we set the scores of unexpressed genes to zero, adopting sophisticated methods to infer cell-type-specific intercellular interactions may refine intercellular CRISPR screens to help identify more specific targets.

Despite these limitations, the intercellular CRISPR screening method identified nine (**Figure 3, Supplementary Table 3**) and 34 interactions (**Figure 4A**) that are targeted by approved drugs and phase III clinical trial drug candidates, respectively. Moreover, our results suggested seven interactions as potential IO targets (**Table 1**). Focusing on interactions instead of single genes alleviates problems arising from the fact that a single gene may have multiple interaction partners. In addition, the novel method proposed in the study is useful because ligands and receptors are more druggable. A recent study suggested that a *PDCD1*-*CD274* bi-specific antibody is more effective than using anti-*PDCD1* and anti-*CD274* antibodies together (54), further supporting the use of intercellular CRISPR screens relative to single screens.

With the aforementioned advantages, intercellular CRISPR screens can be used to evaluate interactions between several cancer types and immune cells such as natural killer cells or macrophages (15, 55). Moreover, the proposed method can be extended to consider multiple immune cell types simultaneously to more precisely model tumor microenvironment.

## 5 Conflict of Interest

NH is a cofounder of KURE.ai and CardiaTec Biosciences and an advisor at Biorelate, Promatix, Standigm, VeraVerse, and Cellaster. All other authors declare that they have no commercial or financial relationships that could be construed as a potential conflict of interest.

## 6 Author Contributions

SY and WH conceptualized the study. SY developed and evaluated the proposed method along with WH under the supervision of NH and DL. All authors contributed to writing, reading, and have approved the manuscript.

## 7 Funding

NH and WH were funded by LifeArc. The work described and publication of this article were supported by the Bio-Synergy Research Project (NRF-2012M3A9C4048758) of the Ministry of Science and ICT through the National Research Foundation.

## 8 Abbreviations

CRISPR: clustered regularly interspaced short palindromic repeats
CTL: cytotoxic T lymphocyte
TNBC: triple-negative breast cancer
IO: immuno-oncology
ATC: anatomical therapeutic chemical
sgRNA: single guide RNA
TIL: tumor-infiltrating lymphocyte
KO: knockout
CTRL: control
diff: differential analysis
exp: expression
comb: combination
FDR: false discovery rate
AUROC: area under the receiver operating characteristic curve

## 9 Acknowledgments

Not applicable.

## Notes

https://go.drugbank.com/releases/5-1-8/downloads/all-full-database

https://ftp.ebi.ac.uk/pub/databases/chembl/ChEMBLdb/latest/

https://ars.els-cdn.com/content/image/1-s2.0-S009286741930844X-mmc1.xlsx

https://ars.els-cdn.com/content/image/1-s2.0-S009286741930844X-mmc2.xlsx

https://staticcontent.springer.com/esm/art%3A10.1038%2Fs41586-020-2746-2/MediaObjects/41586_2020_2746_MOESM6_ESM.zip

https://asrhou.github.io/NATMI/#s2

https://www.cancerresearch.org/scientists/immuno-oncology-landscape

