## Supplementary Figures for "Intercellular CRISPR Screens Enhance the Discovery of Cancer Immunotherapy Targets"

Supplementary Material

### 1 Supplementary Figures and Tables

#### 1.1 Supplementary Figures

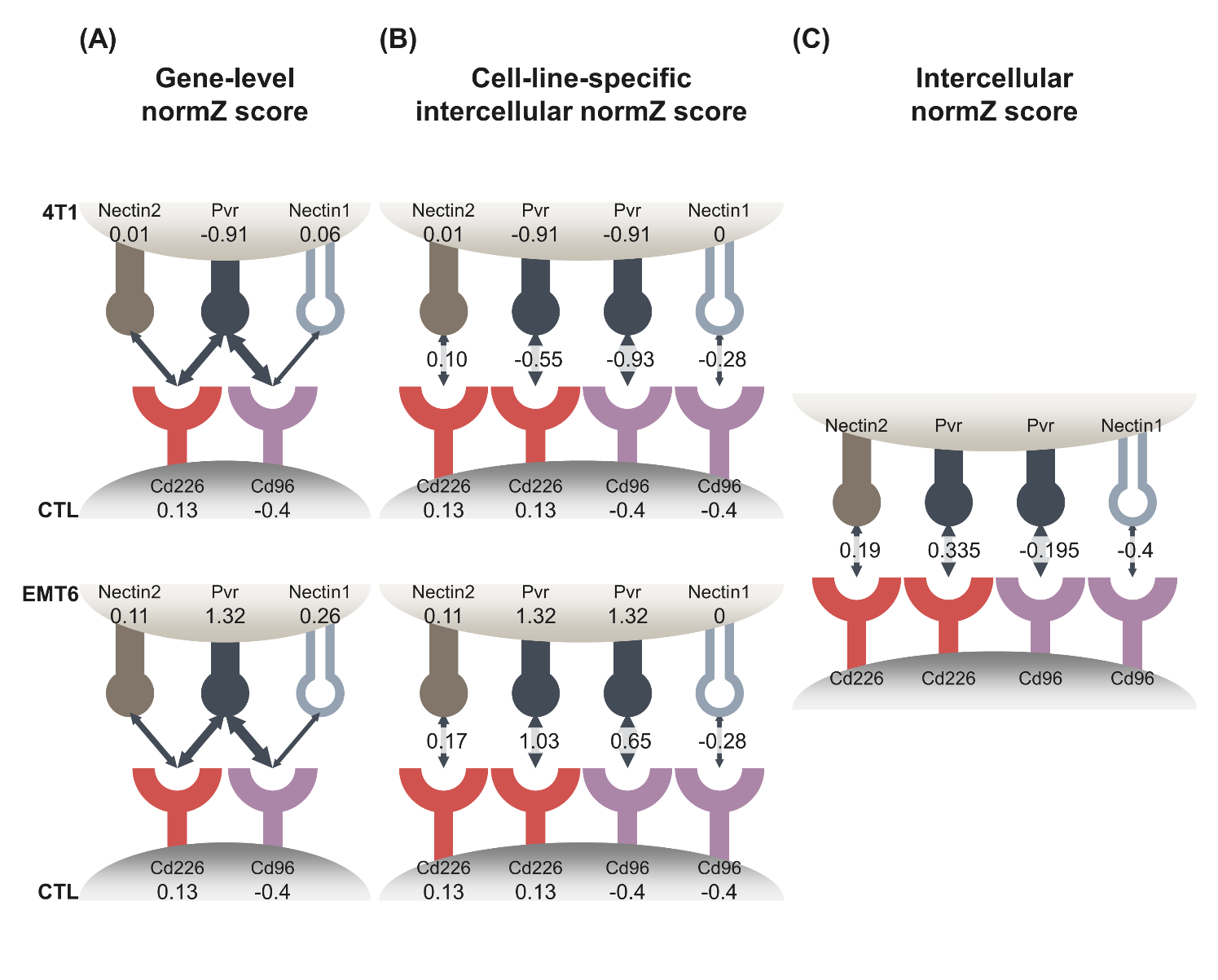

**Supplementary Figure 1.** An example for the calculation of intercellular normZ scores. **(A)** Gene-level normZ scores for CTLs and two TNBC cell lines, 4T1 and EMT6, were calculated by applying the drugZ algorithm to the normalized read count matrices. **(B)** Gene-level normZ scores of Nectin1 in 4T1 and EMT6 were set to zero because Nectin1 was not expressed in both TNBC cell lines. The cell-line-specific intercellular normZ score was calculated by summing the gene-level normZ scores of the ligand and the receptor and dividing the sum by square root of two. **(C)** The final intercellular normZ score was calculated by summing two cell-line-specific intercellular normZ scores and dividing the sum by the square root of two. CTL: cytotoxic T lymphocyte.

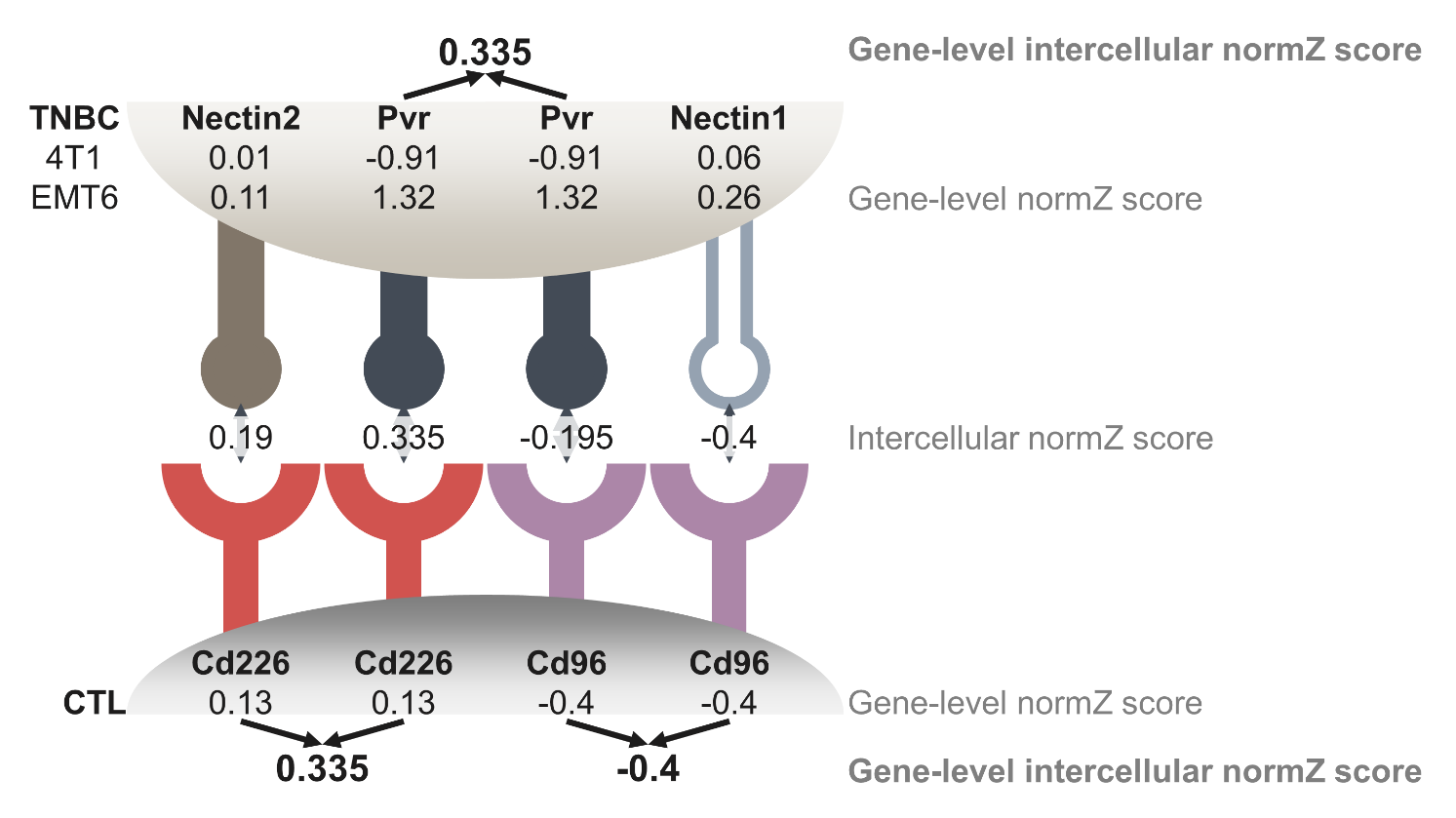

Supplementary Figure 2. An example for aggregating intercellular normZ scores into a gene-level score. Gene-level normZ scores of genes are indicated below the gene symbols. Intercellular normZ scores of interactions are indicated between ligands and receptors. Gene-level intercellular normZ scores for a gene in CTL/TNBC cells were obtained by taking the intercellular normZ score with the highest absolute value. TNBC: triple-negative breast cancer, CTL: cytotoxic T lymphocyte.

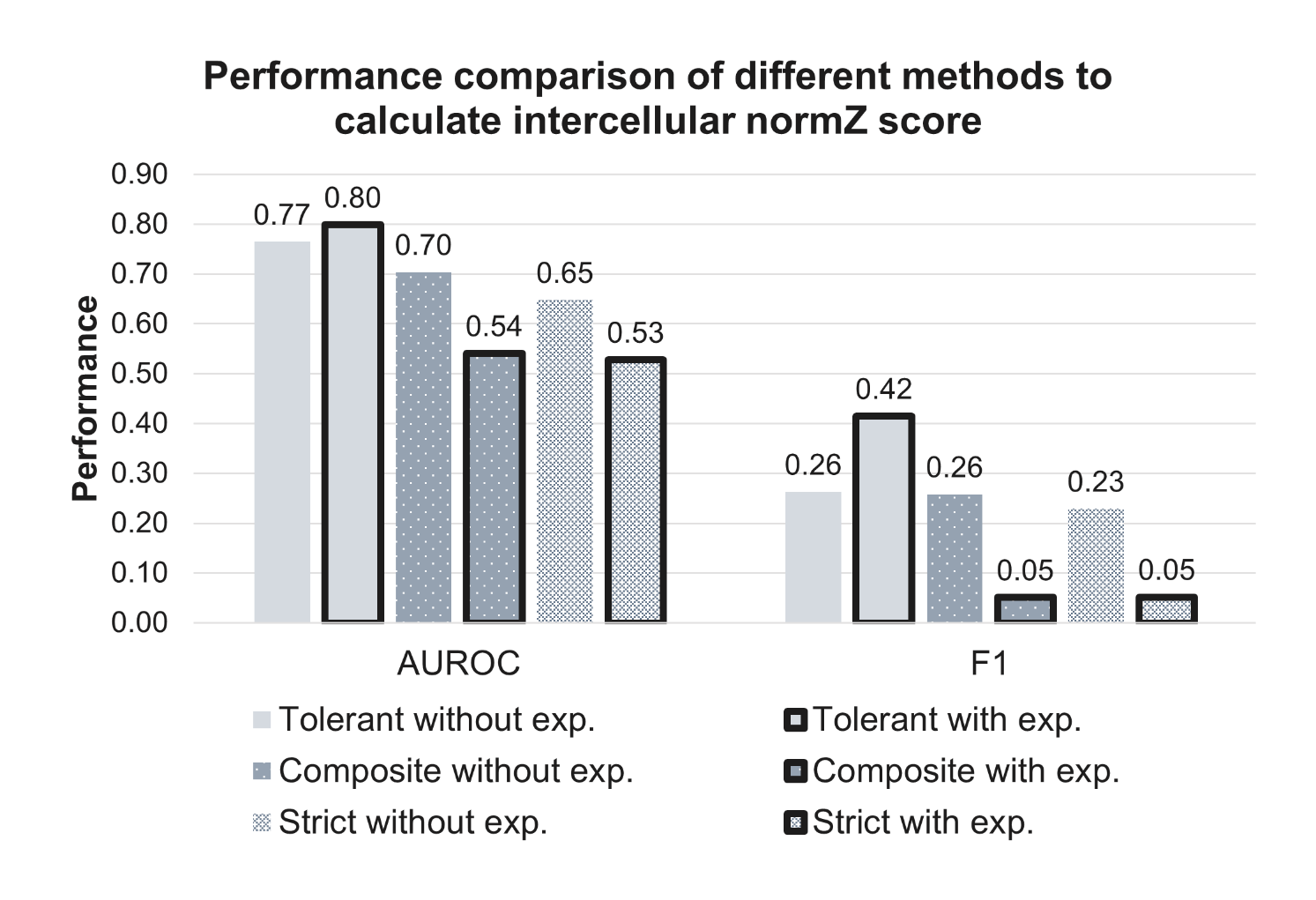

Supplementary Figure 3. Performance comparison of different methods to calculate intercellular normZ scores. ‘Tolerant’ without expression data achieved the highest AUROC and F1 scores. AUROC: area under the receiver operating characteristic curve, exp.: expression.

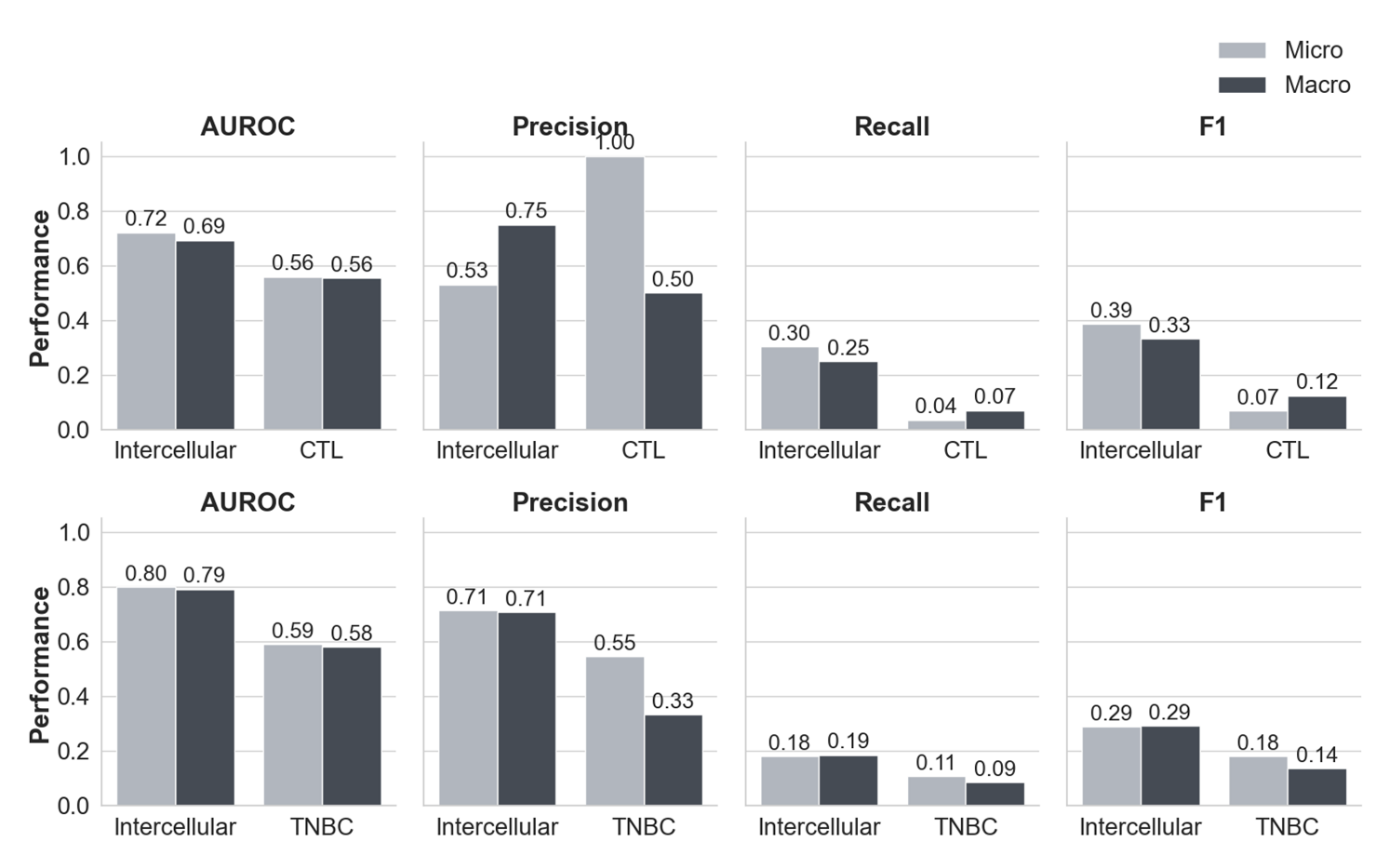

Supplementary Figure 4. The performance comparison between single and gene-level intercellular CRISPR screens. The gene-level intercellular CRISPR screen outperforms single CRISPR screens except for the micro-averaged precision of the CTL CRISPR screen. AUROC: area under the receiver operating characteristic curve, CTL: cytotoxic T lymphocyte, TNBC: triple-negative breast cancer.

#### 1.2 Supplementary Tables

**Supplementary Table 1**. The statistics of approved drugs and their targets. IO: immuno-oncology.

**Supplementary Table 2**. Intercellular normZ scores and corresponding FDRs for all intercellular interactions. FDR: false discovery rate, CTL: cytotoxic T lymphocyte, TNBC: triple-negative breast cancer.

**Supplementary Table 3**. Gold standard dataset of 38 intercellular interactions targeted by approved drugs or phase III clinical trial drug candidates for immunotherapy. TNBC: triple-negative breast cancer, CTL: cytotoxic T lymphocyte.
